# Iris Crypts Could Reduce the Chance of Angle Closure: A Computational Biomechanics Study Derived from Clinical and Human Iris Data

**DOI:** 10.1101/2024.12.30.630843

**Authors:** Royston K.Y. Tan, Tin A. Tun, Fabian A. Braeu, Shamira A. Perera, Michaël J.A. Girard

**Affiliations:** Singapore Eye Research Institute, Singapore National Eye Centre, Singapore; Duke-NUS Medical School, Singapore; Singapore National Eye Centre, Singapore; Department of Biomedical Engineering, Georgia Institute of Technology and Emory University, Atlanta, United States; Department of Ophthalmology, Emory University School of Medicine, Atlanta, United States; Emory Empathetic AI for Health Institute, Emory University, Atlanta, GA, USA

**Keywords:** Iris crypts, permeability, pupil dilation, angle closure

## Abstract

**Purpose:** To investigate the effect of crypts during pupil constriction and dilation on the potential for angle closure by performing finite element analysis using clinical and experimental data on human tissues.

**Methods:** A computational model was developed to determine the influence of small crypts (surface area of ∼0.015 mm^2^) and large crypts (surface area of 0.300 mm^2^) on the anterior chamber angles during pupil dilation. The model needed permeability data from human subjects, hence 21 enucleated human eyes (72 hours post-mortem) were procured and subjected to a flow set-up previously reported. Finally, 66 subjects were recruited to measure pupil constriction and dilation levels from optical coherence tomography (OCT) videos.

**Results:** The hydraulic permeability of the human iris stroma was determined to be 2.55 ± 1.93 × 10^-5^ mm^2^/Pa*·*s. The average iris constriction and dilation durations were 0.710 ± 0.213 seconds and 1.24 ± 0.401 seconds, respectively, with pupil diameter changes of 1.16 ± 0.39 mm and 0.75 ± 0.27 mm, respectively. The computational models had starting anterior chamber angles of 51.30° and final anterior chamber angles of 26.81° once steady state has been reached. In an extreme case with decreased anterior border layer (ABL) permeability, the anterior chamber angle narrowed to 12.37°, but the presence of crypts kept the angle above 20.36°, highlighting the potential of crypts in preventing angle closure.

**Conclusions:** Our findings on the biomechanics of crypts in the iris may drive the development of novel treatments by altering ABL morphology, providing an alternative bypass for angle closure prevention in high-risk patients.

## Introduction

Angle closure glaucoma (ACG) occurs when the anterior chamber drainage angle becomes obstructed, leading to a rise in intraocular pressure (IOP), which damages the cells in the optic nerve leading to vision loss.^1, 2^ The underlying mechanism of angle closure involves restricted trabecular aqueous humor outflow, which can be influenced by both anatomical and dynamic factors.^3, 4^ Therefore it is crucial to explore the factors and processes within the anterior chamber that contributes to this phenomenon.

Movement of the iris during pupil dilation differs for each individual, and the presence of surface features such as crypts had been hypothesized to play a part in angle closure. The morphology of the anterior border layer (ABL) is a dense matrix, comprised of discontinuous collagen fibrils, fibroblasts and melanocytes.^5^ Underneath lies the iris stroma with a similar composition, albeit with a much lower density.^6^ Aqueous humor freely moves within the pores of these layers and makes up to 40% of the stroma layer.^6, 7^ This combination of tissue, along with the dilator muscle posterior to the stroma and sphincter muscle at the iris margin, forms a unique composition with physiological movements greater than most other tissues in the human body. During pupil dilation, the pupil diameter in dark conditions could double when transitioning from light conditions, and likewise, the tissue at the iris margin stretches to twice the length at the iris margin. This causes a large displacement of aqueous humor in a relatively short duration,^8, 9^ which could be the basis for angle closure to occur from excessive bowing when the fluid shift is abnormally slow.

Investigating the various contributing factors for angle closure includes methods such as optical coherence tomography (OCT)^10^ and ultrasound biomicroscopy (UBM)^11^ to obtain images or videos of the anterior segment. Subsequently, a non-invasive method to observe various scenarios *in vivo* is to apply biomechanical principles in computational simulations, such as finite element modeling of recreated anterior chambers.^12–15^ Previously, we investigated six parameters (anterior chamber depth and width, iris convexity, thickness, stiffness and Poisson’s ratio) and found anterior chamber interactions to be highly complex, with each parameter disproportionately contributing to narrowing of the anterior chamber angle.^16^ However, movements in the anterior chamber occur constantly from aqueous flows and iris accommodative muscle actions. Crypts, which are discontinuities of the anterior border layer, could offer alternative pathways for aqueous humor to escape,^17, 18^ leading to smaller dilated iris volumes. In fact, Tun et al.^19^ found that higher crypt grade (i.e., size) was associated with a smaller iris volume and a less convex iris. Similarly, Chua et al.^18^ found that a high crypt grade was associated with a smaller iris volume and greater reduction of iris volume on pupil dilation. However, research on crypts is scarce and the relationship between crypt location and sizes on the anterior chamber angles remain unavailable. Therefore, the aim of this study is to re-create the *in vivo* conditions of the anterior chamber to investigate the effect of crypts in physiological and angle closure scenarios. Specifically, we aimed to determine (1) the permeability of the human iris stroma using *ex vivo* tissues, (2) the speed of the iris during pupil constriction and dilation, and (3) effect of crypts *in vivo* by performing transient computational analysis using clinical and experimental data.

## Methods

This study consisted of two investigational components to determine data that are unavailable in existing literature: permeability of the human iris stroma and the time taken for pupil dilation. These data were crucial for creating a realistic and complex computational model that can closely reflect the behavior of the iris *in vivo*.

### Permeability of the human iris stroma

Twenty-one enucleated healthy Caucasian human eyes (9 male, 12 female, aged 64.3 ± 13.1) were purchased from Saving Sights (De-identified whole globes, Saving Sights, Kansas City, MO, USA). The human eyes were enucleated and placed in a moist chamber at approximately 0°C to 4°C within 12 hours post-mortem. The shipment was kept cold in ice and delivered to the Singapore Eye Research Institute (Singapore National Eye Centre, Singapore) within 72 hours post-enucleation.

The procedure for separating the stroma and dilator tissues was similar to our previous study, which was validated through histological analysis. First, for each eye, the entire iris was isolated from the human globe, and the iris pigmented epithelium was gently scraped off. Next, a discontinuous cut was made to the iris. Using a combination of tying forceps (2-500-2E, Duckworth & Kent, Hertfordshire, UK), jewellers forceps (2-900E, Duckworth & Kent, Hertfordshire, UK) and Chihara conjunctival forceps (2-500-4E, Duckworth & Kent, Hertfordshire, UK), the dilator was removed in small pieces and peeled away from the stroma outward radially up to the sphincter, where the tissue was trimmed using a pair of vannas scissors (1-111, Duckworth & Kent, Hertfordshire, UK). After tissue isolation, the samples were stored in phosphate-buffered saline (PBS).

The isolated stroma samples (*n* = 21) were subjected to a flow experiment to assess their permeability. The principle of the permeability experiment was similar to that described in our previous study using Darcy’s law.^20, 21^ The hydraulic permeability is defined by the permeability of the tissue divided by the viscosity of the fluid, which in this case is PBS:

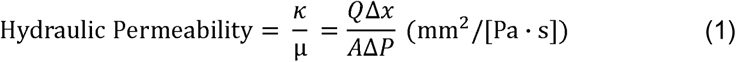

where *K* is the permeability (mm^2^), *μ* is the dynamic viscosity of the fluid (Pa*·*s), *Q* is the flow rate (mm^3^/s), Δ*x* is the thickness of the tissue (mm), *A* is the surface area of flow (mm^2^) and Δ*P* is the pressure drop across the tissue in the direction of flow (Pa). For our experiments, the hydraulic permeability of the iris stroma is determined by setting a specific pressure difference and measuring the resulting flow rate. This setup is identical to our previous study, except for the custom 3D-printed holder used to position the stroma tissue. This area was increased to reduce variance caused by setup misalignments.

### Investigating human iris constriction and dilation duration

The pupillary light response has been extensively studied,^22, 23^ but data on the duration of iris constriction and dilation is less readily available. Hence, a total of 66 patients of Indian and Chinese ethnicities with angle closure, aged 60.2 ± 8.7, were recruited for this study at the Singapore Eye Research Institute, Singapore. Prior informed consent was obtained for all patients. The study was conducted following the tenets of the World Medical Association’s Declaration of Helsinki and received ethics approval from the SingHealth Centralized Institutional review board. All subjects underwent anterior segment imaging using the swept-source OCT (SS-1000 CASIA, Tomey Corporation, Nagoya, Japan) while positioned in the primary gaze position. The OCT imaging was performed with the ‘angle analysis’ protocol in the video mode setting. The anterior segment scan covered 0° to 180°, from limbus to limbus, with an acquisition time of 0.125 seconds per line, and the video was captured at 8 frames per second. The recording of the OCT video started one minute after dark adaptation using a standard protocol, with the light intensity approximately 20 lux measured by a light meter (Studio Deluxe II L-398, Sekonic, Japan). Subsequently, a light source (approximately 1700 lux) was used to flash the subject’s eye from the temporal side, 150° off-axis, to capture the transition from dark to light. The light was turned off after approximately two seconds to capture the transition from light to dark. If motion artifacts were observed, the procedure was repeated, but no more than three times to prevent iris muscle fatigue. Each OCT B-scan was 16 mm in width and 6 mm in depth.

The videos were processed using Adobe Premiere Pro (v25.0, Adobe Inc., San Jose, CA, USA) after exporting the uncompressed data from the CASIA OCT. The start and end points for both constriction and dilation were identified by evaluating the B-scans frame-by-frame to measure the constriction and dilation durations, and the pupil diameters were evaluated by measuring the horizontal opening between the iris margins. Since the level of pupil dilation is less in angle closure subjects,^24^ only the duration would be used for the finite element simulation, but with a larger prescribed dilation matching healthy subjects.^24^

### Finite element simulation

#### 3D Model

The anterior chamber model consisted of the cornea, sclera, lens, iris (consisting of the ABL, stroma, sphincter muscle and dilator muscle), and ciliary body. The model, created with SolidWorks (2023, Dassault Systèmes, Vélizy-Villacoublay, France), was designed based on average healthy literature values: cornea thickness of 0.50 mm,^25, 26^ cornea radius of curvature of 7.50 mm,^27^ sclera radius of curvature of 12.00 mm,^28^ anterior chamber depth (ACD) of 3.00 mm,^29–32^ anterior chamber width (ACW) of 11.50 mm^32–34^ and only the anterior lens of radii of 1.5 mm and 4.75 mm^35, 36^ (**Figure 1A**). The cornea and sclera were combined as a single tissue since the stiffness of the tissues was similar and far greater than the iris (essentially a rigid body). The trabecular meshwork region was also combined with the cornea and sclera, and was angled 15° from the cornea at a small curvature. The lens, intended to be a rigid body during iris sliding in pupil dilation, was simplified as a shell with a thickness of 0.10 mm.

**Figure 1.**
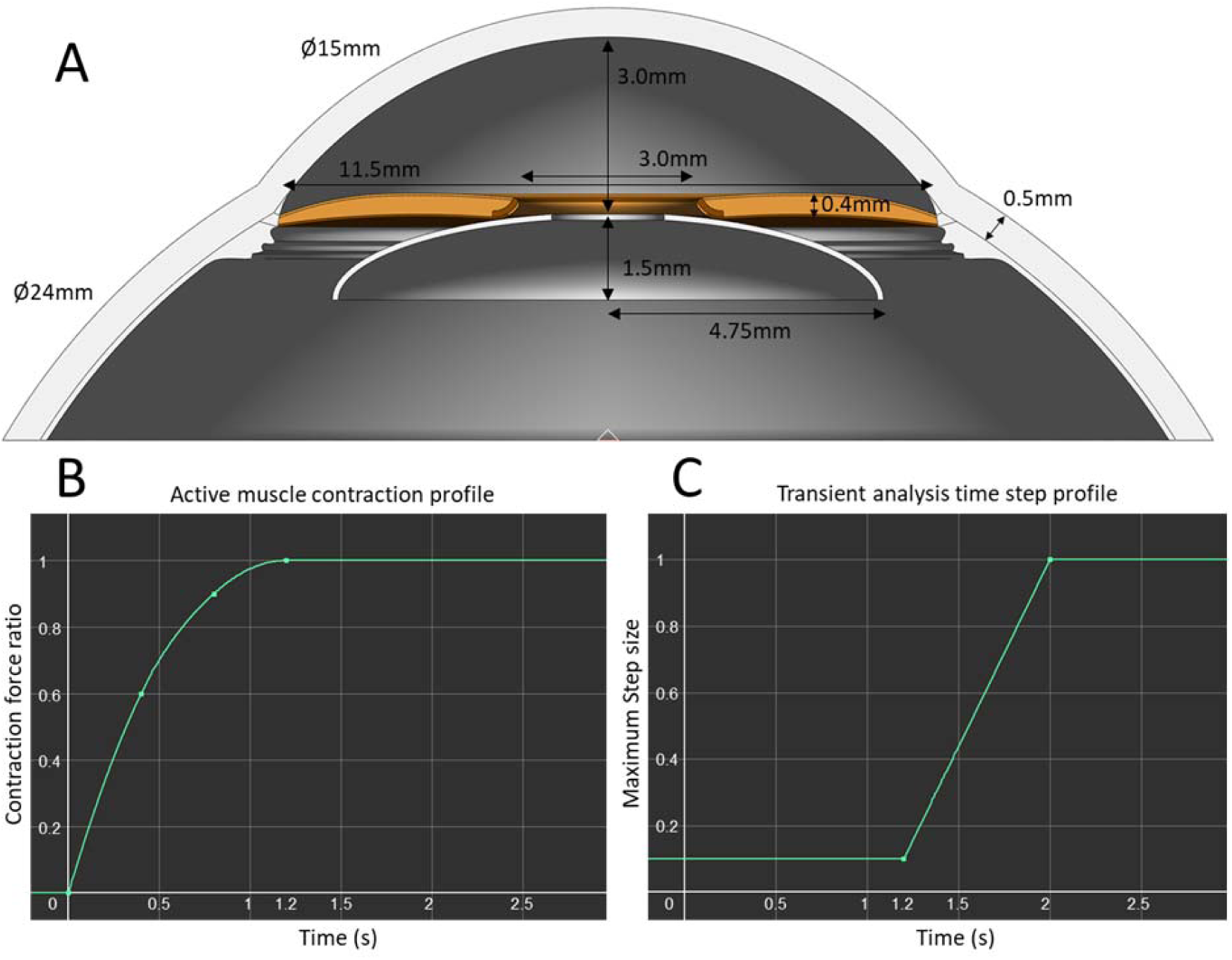
**A.** Anterior chamber parameters for the computational model. The model parameters consisted of **B.** dilator muscle contraction force of 94 kPa for 2.5 mm pupil dilation in 1.2 seconds, using **C.** transient analysis with a custom step size profile.

The iris was attached to the sclera at the root, which was which was consistently set at 0.20 mm thickness for all cases. Iris tissue thickness was defined as distance from the iris margin to the mid-periphery, and it decreased from 0.40 mm at the mid-periphery to the iris root. The iris tissue had a convexity of 0.10 mm, was rounded at the margin, and had a pupil opening of 3.00 mm. The iris contained the sphincter muscle with a thickness of 0.10 mm, located 0.04 mm away from the posterior surface. The dilator muscle was located posterior to the iris, with a thickness of 0.04 mm, spanning from the mid-portion of the sphincter muscle to the iris root. The ABL, spanning the entire anterior surface up to the sphincter muscle, had a thickness of 0.04 mm. The ciliary body was included for illustration purposes but did not influence the simulation results.

#### Mesh and convergence

The 3D model was exported from SolidWorks as a STEP (AP203) file and imported into Abaqus FEA (2021, Dassault Systèmes, Vélizy-Villacoublay, France) for meshing. Eight-node hexahedral elements were used for each model to capture the geometry accurately. A convergence study was performed on an average model with mesh sizes of 6,800, 23,730, 53,120, and 344,960. The convergence test showed that the result for the selected mesh density differed < 0.8% of the result from the most refined mesh, and therefore, the selected mesh (approximately 23,730) was deemed numerically acceptable.

#### Biomechanical parameters

The finite element model contained six parts, each with the following material parameters: The corneo-scleral shell was modeled using an incompressible neo-Hookean formulation, with a Young’s modulus of 500 kPa.^37^ The lens was modeled as a rigid body and was fixed for all scenarios. The sphincter and dilator muscles were described as incompressible neo-Hookean materials, with a Young’s modulus of 40 kPa assigned to each. To induce pupil dilation, the dilator muscle was also described with a uniaxial muscle contraction force of 94 kPa (**Figure 1B**) to dilate the pupil by approximately 2.5 mm. The direction of contraction was along the radial direction of the iris, specified using the element’s local coordinates. Finally, the ABL and iris stroma were described as a biphasic material consisting of a solid neo-Hookean material with a stiffness of 10 kPa and a fluid component with solid fractions of 0.9 and 0.5, respectively. To mimic the volume changes during pupil dilation, the ABL and iris stroma were assigned Poisson’s ratios of 0.3 and 0.1, respectively. Notably, the neo-Hookean formulation was selected in this study primarily for its stability and accuracy when performing large deformations in FEBio.

Using results from the permeability experiments, we assumed that the permeability of the stroma was approximately two orders of magnitude larger than that of the ABL. We then used the harmonic-average permeability to calculate that of the individual layers. Increased cross-linking and changes in tissue composition as part of aging are likely to decrease the permeability of the iris tissues,^38, 39^ with various studies showing a marked decrease in volume change between normal subjects and those with angle closure.^8, 9^ Therefore, permeabilities (µ) of the ABL and stroma were varied to investigate the effect of crypts in various possible transient scenarios: (1) physiological state where µ_ABL_ = 3 × 10^-6^ mm^2^/Pa*·*s and µ_stroma_ = 3 × 10^-4^ mm^2^/Pa*·*s, (2) reduced stroma permeability at 1×µ_ABL_ and 0.1×µ_stroma_, (3) reduced ABL permeability at 0.1×µ_ABL_ and 1×µ_stroma_, and (4) reduced stroma and ABL permeability at 0.1×µ_ABL_ and 0.1×µ_stroma_. The steady state was determined by using ABL and stroma permeabilities at 1 mm^2^/Pa*·*s.

#### Crypts

Images of donor irides were captured and recreated in the computational model with either small or large crypt sizes (**Figure 2A and 2B**). The small crypts had surface areas of 0.015 mm^2^ and the large crypts had surface areas of 0.300 mm^2^, with three crypts present on each quadrant model situated either towards the central region or peripheral region of the iris at approximately 2.7 mm and 4.4 mm pupil radii respectively (**Figure 2C and 2D**).

**Figure 2.**
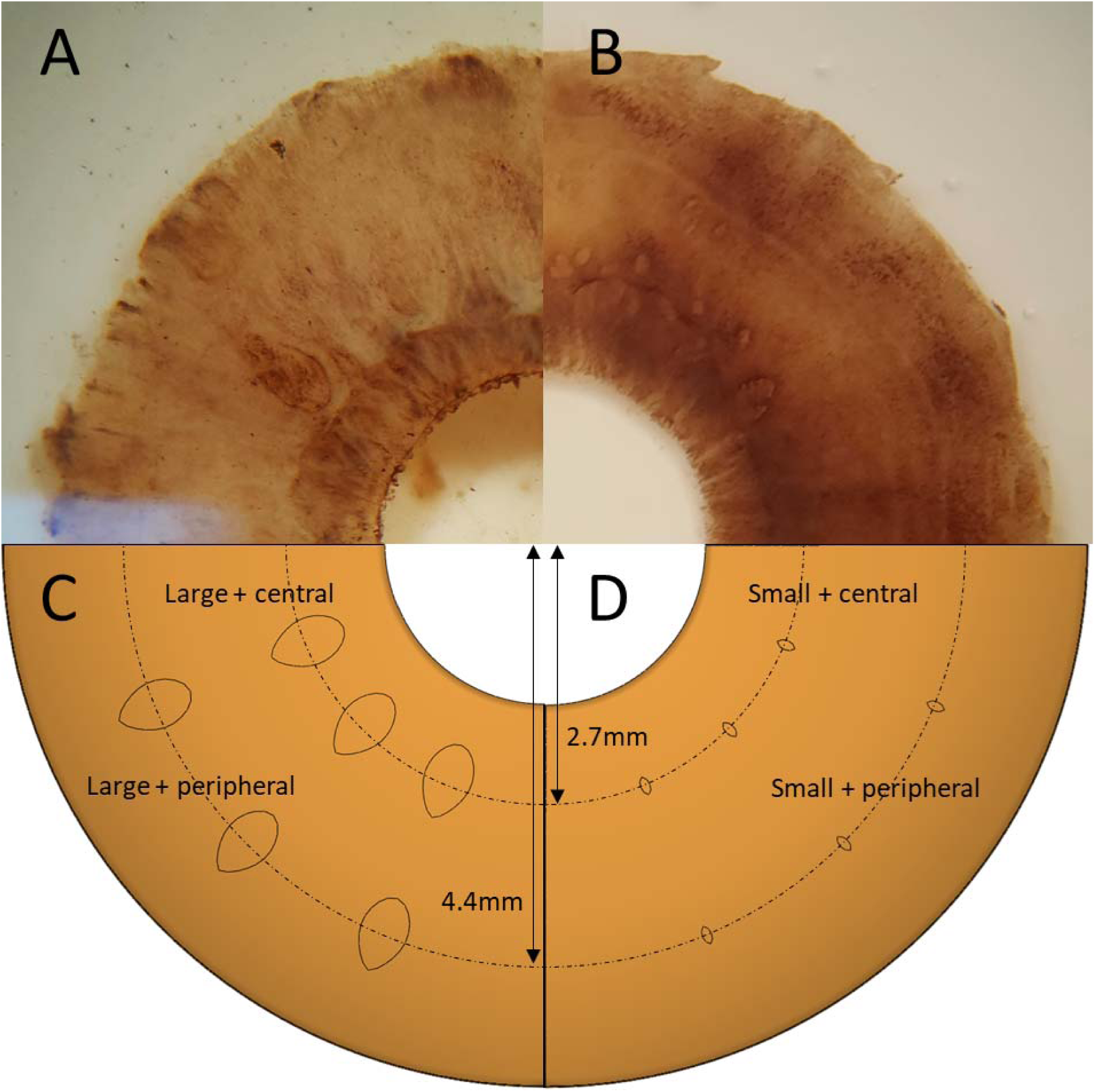
Anterior images of the human iris with **A.** large crypts and **B.** small crypts. The simulation models contained either **C.** 3 large crypts or **D.** 3 small crypts, located at either the central or peripheral locations at approximately 2.7 mm and 4.4 mm radii.

#### Boundary conditions

Constraints along the axisymmetric planes of the quadrant anterior chamber model were applied, and a zero-friction contact interface was assumed between the iris and lens, as well as between the iris and the corneal endothelium surface. The lens was fixed, and so was the outer boundary of the corneo-scleral shell.

#### Finite element processing

All finite element (FE) models were solved using FEBio (FEBio Studio v1.8.2, University of Utah, UT, USA), a non-linear FE solver designed for biomechanical studies. Transient analysis was conducted with dilator muscle contraction lasting 1.2 seconds, and subsequently held for the next 8.8 seconds for a total of 10 seconds at appropriate step sizes (**Figure 1C**) for the model to reach equilibrium. The anterior chamber angles were determined using the coordinates of the nodes at the scleral spur and its adjacent nodes to calculate the angles. We reported the anterior chamber angles at the minimum and after 1.2 seconds of contraction.

## Results

### Permeability of the human iris stroma

The 3D tissue holder area was evaluated using OCT and determined to be 0.538 mm^2^. The average thickness of the stroma tissue samples was 0.210 ± 0.0523 mm. The PBS flow through the stroma samples was found to be slow and laminar at 0.123 ± 0.094 mm^3^/s, with a Reynold’s number < 10, thus satisfying the Darcy’s law approximation. The hydraulic permeability of the human iris stroma was determined to be 2.55 ± 1.93 × 10^-5^ mm^2^/Pa*·*s, with a total of 21 samples evaluated. Hence, the value used to calculate the harmonic-average permeability for the finite element study is 2.5 × 10^-5^ mm^2^/Pa*·*s.

### Investigating human iris constriction and dilation speeds

From the 61 samples evaluated, the average iris constriction and dilation durations were 0.710 ± 0.213 seconds and 1.24 ± 0.401 seconds, respectively. Constriction occurred as the pupil diameter decreased from 3.53 ± 0.60 mm to 2.37 ± 0.38 mm, with the average constriction being 1.16 ± 0.39 mm. Dilation occurred as the pupil diameter increased from 2.39 ± 0.38 mm to 3.14 ± 0.45 mm, with the average dilation being 0.75 ± 0.27 mm. Hence, the approximate duration used for dilation in the finite element model was 1.2 seconds.

### Finite element model

The starting anterior chamber angle for all models was 51.30°, which narrowed to a minimum (depending on the scenario) after approximately 2.5 mm of pupil dilation but then widened to a final angle of 26.81° once steady state was reached (**Table 1**).

**Table 1.** Changes in anterior chamber angles (Δ∠) from the initial non-dilated (ACA_i_) state at *t* = 0 to immediately after dilator muscle activity (ACA_1.2s_) at *t* = 1.2s. The minimum anterior chamber angle (∠_minimum_) occurs in this duration, determined using the transient analysis. 1×µ_ABL_ = 3 × 10^-6^ mm^2^/Pa*·*s, 0.1×µ_ABL_ = 3 × 10^-7^ mm^2^/Pa*·*s, 1×µ_stroma_ = 3 × 10^-4^ mm^2^/Pa*·*s and 0.1×µ_stroma_ = 3 × 10^-5^ mm^2^/Pa*·*s.

| **Iris Feature** | **Permeabilities** | **$ACA_i$ (°)** | **$\Delta\angle$ (°)** | **$ACA_{1.2s}$ (°)** | **$\angle_{\text{minimum}}$ (°)** |
| --- | --- | --- | --- | --- | --- |
| NA | Steady state | 51.30 | 24.49 | 26.81 | 26.81 |
| No crypts | $1 \times \mu_{\text{ABL}}$ and $1 \times \mu_{\text{stroma}}$ | 51.30 | 27.18 | 24.12 | 23.47 |
| | $1 \times \mu_{\text{ABL}}$ and $0.1 \times \mu_{\text{stroma}}$ | 51.30 | 28.55 | 22.75 | 21.59 |
| | $0.1 \times \mu_{\text{ABL}}$ and $1 \times \mu_{\text{stroma}}$ | 51.30 | 38.37 | 12.93 | 12.37 |
| | $0.1 \times \mu_{\text{ABL}}$ and $0.1 \times \mu_{\text{stroma}}$ | 51.30 | 48.29 | 3.00 | 2.03 |
| 3 small crypts<br>Central region | $1 \times \mu_{\text{ABL}}$ and $1 \times \mu_{\text{stroma}}$ | 51.30 | 26.89 | 24.41 | 23.76 |
| | $1 \times \mu_{\text{ABL}}$ and $0.1 \times \mu_{\text{stroma}}$ | 51.30 | 28.54 | 22.76 | 21.61 |
| | $0.1 \times \mu_{\text{ABL}}$ and $1 \times \mu_{\text{stroma}}$ | 51.30 | 36.53 | 14.77 | 13.78 |
| | $0.1 \times \mu_{\text{ABL}}$ and $0.1 \times \mu_{\text{stroma}}$ | 51.30 | 48.26 | 3.04 | 2.03 |
| 3 large crypts<br>Central region | $1 \times \mu_{\text{ABL}}$ and $1 \times \mu_{\text{stroma}}$ | 51.30 | 26.57 | 24.73 | 24.10 |
| | $1 \times \mu_{\text{ABL}}$ and $0.1 \times \mu_{\text{stroma}}$ | 51.30 | 28.62 | 22.68 | 21.51 |
| | $0.1 \times \mu_{\text{ABL}}$ and $1 \times \mu_{\text{stroma}}$ | 51.30 | 34.16 | 17.14 | 15.58 |
| | $0.1 \times \mu_{\text{ABL}}$ and $0.1 \times \mu_{\text{stroma}}$ | 51.30 | 48.18 | 3.12 | 2.11 |
| 3 small crypts<br>Peripheral<br>region | $1 \times \mu_{\text{ABL}}$ and $1 \times \mu_{\text{stroma}}$ | 51.30 | 26.55 | 24.75 | 24.15 |
| | $1 \times \mu_{\text{ABL}}$ and $0.1 \times \mu_{\text{stroma}}$ | 51.30 | 28.55 | 22.75 | 21.60 |
| | $0.1 \times \mu_{\text{ABL}}$ and $1 \times \mu_{\text{stroma}}$ | 51.30 | 33.71 | 17.59 | 16.40 |
| | $0.1 \times \mu_{\text{ABL}}$ and $0.1 \times \mu_{\text{stroma}}$ | 51.30 | 48.12 | 3.18 | 2.21 |
| 3 large crypts<br>Peripheral<br>region | $1 \times \mu_{\text{ABL}}$ and $1 \times \mu_{\text{stroma}}$ | 51.30 | 25.97 | 25.33 | 24.87 |
| | $1 \times \mu_{\text{ABL}}$ and $0.1 \times \mu_{\text{stroma}}$ | 51.30 | 28.56 | 22.74 | 21.58 |
| | $0.1 \times \mu_{\text{ABL}}$ and $1 \times \mu_{\text{stroma}}$ | 51.30 | 29.65 | 21.65 | 20.36 |
| | $0.1 \times \mu_{\text{ABL}}$ and $0.1 \times \mu_{\text{stroma}}$ | 51.30 | 47.70 | 3.60 | 2.56 |

In the absence of crypts, the anterior chamber angles for the four scenarios were 24.12°, 22.75°, 12.93°, and 3.00° immediately after dilation at 1.2 seconds, with minimum values of 23.47°, 21.59°, 12.37°, and 2.03°, respectively.

For small crypts located at the central region, the angles were 24.41°, 22.76°, 14.77°, and 3.04° immediately after dilation at 1.2 seconds, with minimum values of 23.76°, 21.61°, 13.78°, and 2.03°, respectively.

For large crypts located at the central region, the angles were 24.73°, 22.68°, 17.14°, and 3.12° immediately after dilation at 1.2 seconds, with minimum values of 24.10°, 21.51°, 15.58°, and 2.11°, respectively.

For small crypts located at the peripheral region, the angles were 24.75°, 22.75°, 17.59°, and 3.18° immediately after dilation at 1.2 seconds, with minimum values of 24.15°, 21.60°, 16.40°, and 2.21°, respectively.

For large crypts located at the peripheral region, the angles were 25.33°, 22.74°, 21.65°, and 3.60° immediately after dilation at 1.2 seconds, with minimum values of 24.87°, 21.58°, 20.36°, and 2.56°, respectively.

## Discussion

This study combined human tissue and clinical data into a computational model to understand how crypts could influence angle closure. The models revealed changes in anterior chamber angles with varying crypt sizes and locations, suggesting that decreased permeability in the iris tissue may lead to angle closure in the absence of crypts.

### The size and location of crypts are equally important

When we examined the physiological scenario where µ_ABL_ = 3 × 10^-6^ mm^2^/Pa*·*s and µ_stroma_ = 3 × 10^-4^ mm^2^/Pa*·*s, the model with no crypts showed the greatest fluctuation in anterior chamber angles, with a minimum angle of 23.47° which increased to 24.12° when *t* = 1.2 seconds. The presence of large peripheral crypts allowed for slightly larger anterior chamber angles, with a minimum angle of 24.87° which increased to 25.33° when *t* = 1.2 seconds. Although the size difference between small and large crypts was about 20 times, the presence of small peripheral crypts had similar effects to large central crypts, with minimum values of 24.15° and 24.10°, respectively, increasing to 24.75° and 24.73° at *t* = 1.2 seconds (**Figure 3A**). Similarly at all other scenarios, having small peripheral crypts fared equal or slightly better than having large central crypts, which indicates that the location of crypts is as important as the size of crypts (**Figure 3B to 3D**). Since the peripheral iris contains more volume (thus losing more volume during dilation), it stands to reason that having peripheral crypts allows for more effective aqueous humor outflow. It is important to note that in these physiological scenarios, the variation of anterior chamber angles is low, and the angles converge within approximately 3 seconds. While there are slight differences in the angle profiles, the average human eye, with or without crypts, is capable of pupil accommodation effectively without risks of angle closure.

**Figure 3.**
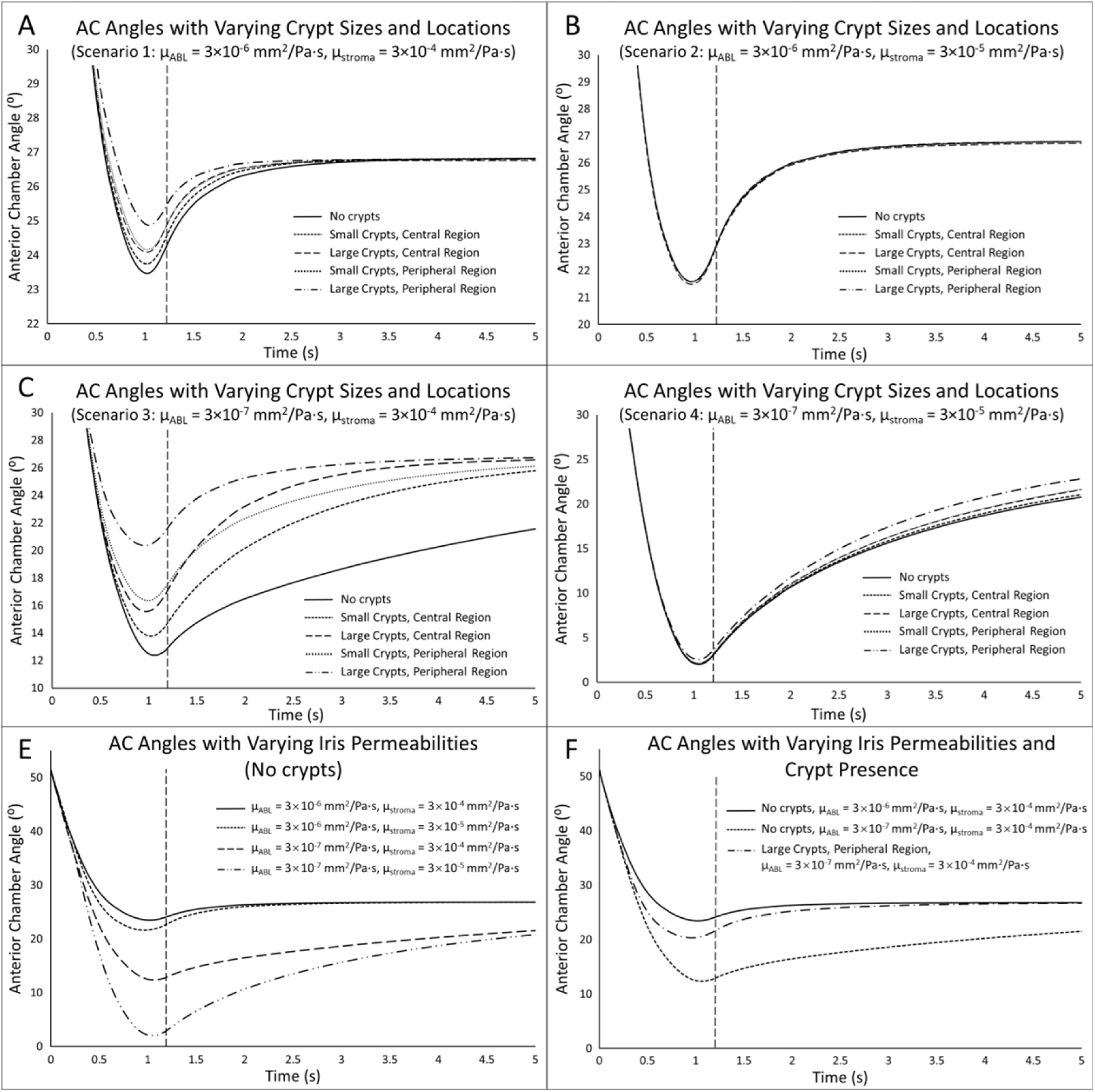
Graphs of anterior chamber (AC) angles with varying crypt sizes and locations in **A.** scenario 1: µ_ABL_ = 3 × 10^-6^ mm^2^/Pa*·*s and µ_stroma_ = 3 × 10^-4^ mm^2^/Pa*·*s, **B.** Scenario 2: µ_ABL_ = 3 × 10^-6^ mm^2^/Pa*·*s and µ_stroma_ = 3 × 10^-5^ mm^2^/Pa*·*s, **C.** µ_ABL_ = 3 × 10^-7^ mm^2^/Pa*·*s and µ_stroma_ = 3 × 10^-4^ mm^2^/Pa*·*s, **D.** Scenario 4: µ_ABL_ = 3 × 10^-7^ mm^2^/Pa*·*s and µ_stroma_ = 3 × 10^-5^ mm^2^/Pa*·*s. **E.** AC angles in the scenario with no crypts at various ABL and stroma permeabilities, and **F.** AC angles showing the effectiveness of large crypts in scenario 3 in preventing narrow angles.

### The presence of crypts could be important in preventing angle closure

The less resistant pathway created by crypts could facilitate more rapid aqueous outflow. When we analyzed the case where no crypts were present (scenario 1), the anterior chamber angle reached equilibrium rapidly under physiological permeability conditions (**Figure 3E**). When stroma permeability decreased (scenario 2), greater fluctuations in the anterior chamber angles were observed. However, when ABL permeability decreased (scenarios 3 and 4), the anterior chamber angle dropped rapidly and failed to reach equilibrium within the same time frame due to reduced aqueous outflow from the iris. In the latter two scenarios, the anterior chamber angles fell below 15°, which could lead to angle closure when accounting for *in vivo* processes such as pupillary block and increased iris convexity caused by posterior chamber pressure differences. However, the presence of crypts has the potential to prevent angle closure. In scenario 3, without crypts, the anterior chamber angle dropped to a minimum of 12.37°, while the presence of large peripheral crypts kept the angle above 20.36°. This also allowed the iris to behave similarly to a healthy iris, reaching equilibrium rapidly (**Figure 3F**). For subjects who are more susceptible to angle closure, i.e., with smaller ACD and ACW and thus smaller anterior chamber angles to begin with, the presence of crypts could very well prevent collapse of the angles.

### The ABL could be the limiting factor in angle closure

The dissection of human globes revealed observable differences reported in literature. The anterior boarder layer is likely to limit aqueous humor flow in angle closure because of its high density. When looking at the anterior view of the iris (**Figure 4A**), the continuous layer is very dense, with non-visible pores covering the area. Freddo had previously reported images of this layer, showing pores (approximately 10–50 µm) visible only under high magnification.^7^ Conversely, the radial fibers of the stroma consisted mainly of fine collagen fibers and iris blood vessels (**Figure 4B**) in a large fraction of extracellular matrix and aqueous humor.^7, 40^ The difference in tissue density translates to a large difference in fluid permeability, which we estimated to be approximately two orders of magnitude. Comparing the simulation scenarios with decreased stroma permeability (**Figure 3A and 3B**), we observed less impact in anterior chamber angles and the duration for the iris tissue to reach a steady state. Conversely, decreased ABL permeability greatly decreases the anterior chamber angles (**Figure 3C and 3D**) and for longer durations. Therefore, an investigation into the surface morphology of iris tissues in patients, specifically ABL pore and crypt sizes, could uncover new correlations with the incidence of angle closure.

**Figure 4.**
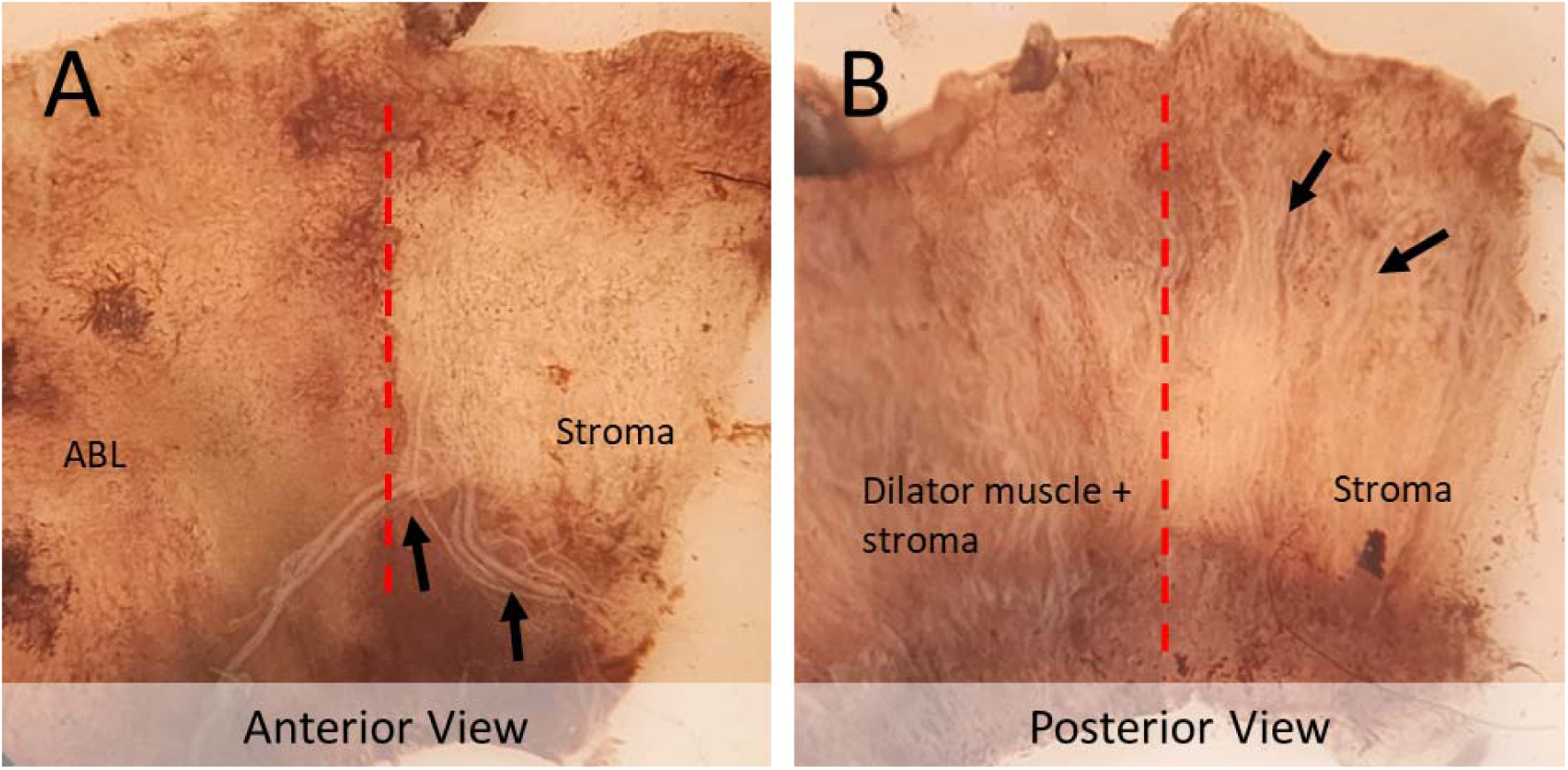
**A.** The anterior and **B.** posterior views of the human iris samples. The areas left of the dotted red lines contain the ABL and dilator muscle respectively, and the areas on the right of the dotted red lines show the visible sparse stroma fibers comprised of mainly collagen strands and iris blood vessels.

**Figure 5.**
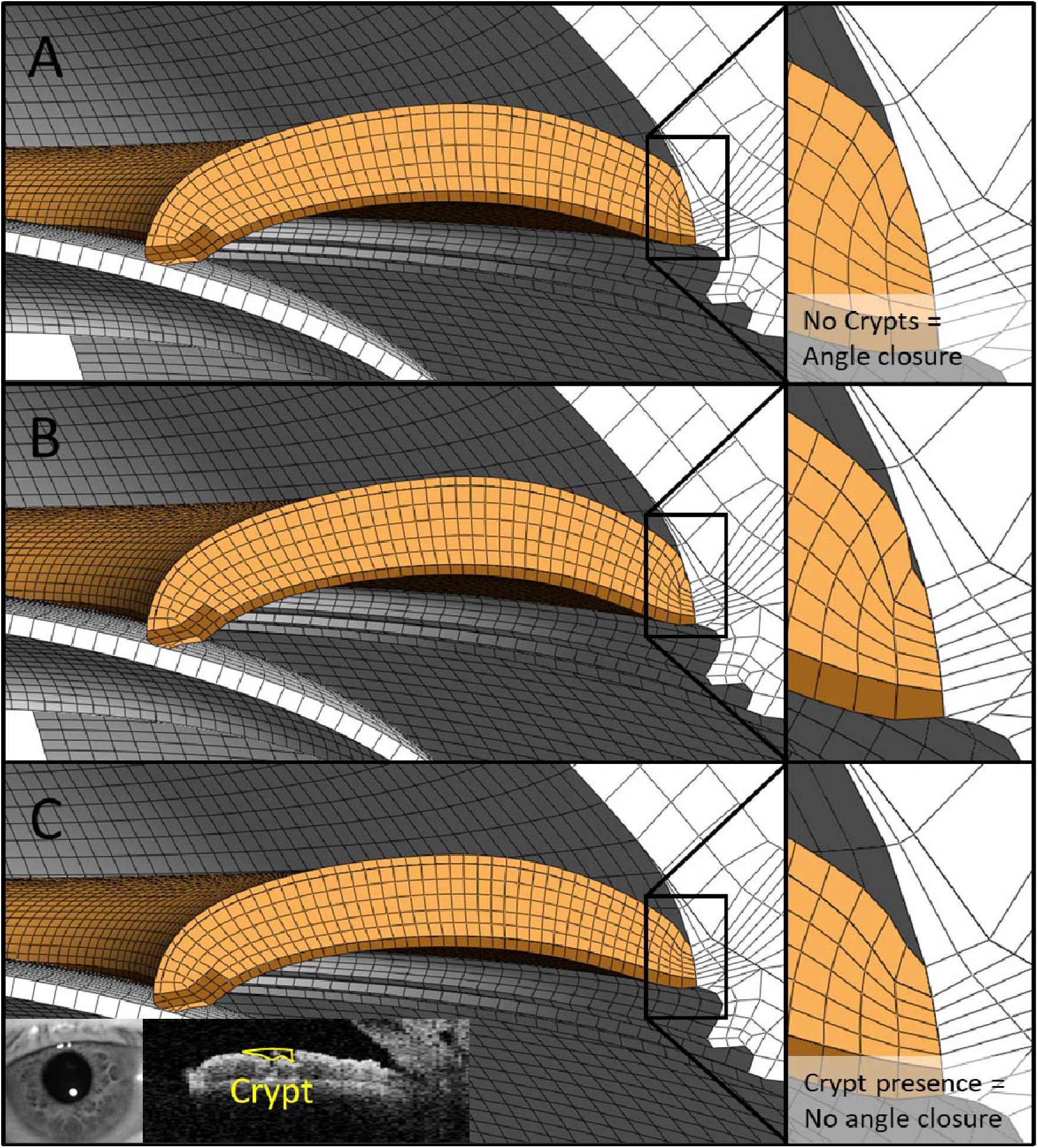
**A.** The case for angle closure when the ABL layer permeability decreases to restrict fluid outflow, when no crypts are present. **B.** An iris (with crypts) may still cause instantaneous angle closure when the iris dilates, but **C.** The presence of crypts could be significant in facilitating aqueous outflow to increase the anterior chamber angle, evident in an OCT scan of a patient with narrow angle and a visible crypt.

### Limitations

In combining experimental, clinical, and computational data, several assumptions were made to reach the conclusions outlined above. First, permeability values were determined with the assumption that ABL permeability is 100 times lower than that of the stroma, based on histological studies and calculated using harmonic average permeability with a thickness ratio of 1:9. Changing this assumption to 200 times, for example, would shift the limiting factor further towards the ABL, and vice versa. This ratio may vary between subjects and warrants further investigation.

Second, the dilation duration was based on a population of older angle closure subjects, including individuals with angle closure suspect, primary angle closure, and primary angle closure glaucoma. This dilation speed (1.24 ± 0.401 seconds) and dilation amount (0.75 ± 0.27 mm) would differ for a younger, healthier population. To compensate for this, we selected a dilation duration of 1.2 seconds for a 2.5 mm dilation, consistent with existing literature.^41^ However, it is also important to recognize that, while constriction is rapid, dilation is not only slower but can take up to a minute for the pupil to fully dilate. Therefore, the constriction profile (**Figure 1B**) is important for reflecting physiological dilation profiles, which can also vary between individuals.

Third, the pupil is constantly undergoing antagonistic muscle accommodation to regulate the amount of light entering the eye. This makes iris biomechanics complex, as the iris muscles are constantly constricting or relaxing.^42^ We simplified iris movement during pupil dilation as being solely due to the constriction of the dilator muscle, with the sphincter muscle considered passive. This means the pupil started from a neutral, relaxed position. However, if the pupil begins from a constricted state, the sphincter muscle should have been assigned a pre-tensile force to shift it from the neutral position before dilation.

Finally, the anterior chamber contains other aqueous humor processes that affect the anterior chamber, such as production from the ciliary body, outflow from the trabecular meshwork,^43, 44^ and the pressure differences between the anterior and posterior chambers of the iris.^45^ These processes are dynamic and interact with iris muscle movement, affecting iris curvature and narrowing the anterior chamber angles. However, they were omitted from the study due to complexity and difficulty in replicating physiological conditions *in vivo*.

## Conclusion

Our findings on the biomechanics of iris crypts offer new insights into potential strategies for preventing angle closure in high-risk patients. By altering the morphology of the anterior border layer (ABL), we may create an alternative pathway for aqueous humor outflow, mitigating the risk of angle closure. To move this concept forward, further exploration of the microscopic structure of the ABL and iris stroma is essential. Understanding these tissue characteristics in greater detail will be critical for developing effective preventive treatments or in risk stratifying subjects with angle closure.

## Acknowledgements

The authors thank the donors of the National Glaucoma Research, a program of the BrightFocus Foundation, for support of this research (G2021010S [MJAG]); the NMRC-LCG grant ‘TAckling & Reducing Glaucoma Blindness with Emerging Technologies (TARGET)’, award ID: MOH-OFLCG21jun-0003 [MJAG]; and the SERI-Lee Foundation grant (LF0622- 03).

